# Primary Evidence of a Donut-Like, Fourth Spatial Dimension in the Brain

**DOI:** 10.1101/072397

**Authors:** James F. Peters, Ebubekir İnan, Arturo Tozzi, Sheela Ramanna

## Abstract

We introduce a novel method for the measurement of information in fMRI neuroimages, *i.e*., nucleus clustering’s Rényi entropy derived from strong proximities in feature-based Voronoï tessellations, *e.g.*, maximal nucleus clustering (MNC). We show how MNC is a novel, fast and inexpensive image-analysis technique, independent from the standard blood-oxygen-level dependent signals, which facilitates the objective detection of hidden temporal patterns of entropy/information in zones of fMRI images generally not taken into account by the subjective standpoint of the observer. In order to evaluate the potential applications of MNC, we looked for the presence of a fourth dimension’s distinctive hallmarks in a temporal sequence of 2D images taken during spontaneous brain activity. Indeed, recent findings suggest that several brain activities, such as mind-wandering and memory retrieval, might take place in the functional space of a four dimensional hypersphere, which is a double donut-like structure undetectable in the usual three dimensions. We found that the Rényi entropy is higher in MNC areas than in the surrounding ones, and that these temporal patterns closely resemble the trajectories predicted by the possible presence of a hypersphere in the brain.

---

In this paper we introduce a novel technique of fMRI images analysis, called computational proximity method, *i.e*., nucleus clustering in Voronoï tessellations (Peters and Inan, 2016a). The images are subdivided in contiguous (without interstice or overlap) polygons, called the “Voronoï polygons”. They yield a density map, called “tessellation”, that makes it possible to make an objective measurement of the polygon areas’ spatial distribution and helps to define “random”, “regular” and “clustered” distributions (Duyckaerts and Godefroy, 2000; Franck and Hart, 2010; Edelsbrunner, 2014). Tessellations have been already used in neuroscience, *i.e.*, to investigate spatial relations and connectivity between neural mosaics in the retina (Mozos et al., 2011) or to evaluate histological cortical sections (Peters at al., 2016). In a Voronoï tessellation of an fMRI image, of particular interest is the presence of maximal nucleus clusters (MNC), *i.e*., zones with the highest number of adjacent polygons (Peters et al., 2016). The MNC clustering approach includes a main feature of level set methods, namely, a nucleus boundary that is embedded in a family of nearby level sets (Saye and Sethian, 2011). MNC reveals regions of the brain, independent from blood-oxygen-level dependent (BOLD) signals, characterized by different gradient orientation and diverse functional dimensions.

To evaluate the power and potentialities of this novel approach, we used it in order to test the brain-hypersphere hypothesis. Indeed, it has been recently hypothesized that brain activity is shaped in the guise of a functional hypersphere, which performs complicated 4D movements called “quaternionic” rotations (Tozzi and Peters, 2016a). They give rise to the so called “Clifford torus”, a closed donut-like structure where mental functions might take place. The torus displays glued trajectories similar to a video game with spaceships in combat: when a spaceship flies off the right edge of the screen, it does not disappear but rather comes back from the left (Weeks, 2002). The human brain exhibits similar behaviour, *i.e.*, the unique ability to connect far-flung events in a single, coherent picture (Atasoy et al., 2016). During brain functions such as memory retrieval and mind-wandering, concepts flow from one state to another and appear to be “glued” together. It has also been recently proposed that the brain, when evaluated in the proper dimension (Kida et al., 2016), is equipped with symmetries in one dimension that disappear (said to be “hidden” or “broken”) in just one dimension lower (Tozzi and Peters, 2016b). A symmetry break occurs when the symmetry is present at one level of observation, but “hidden” at another level: it suggests that a 4D hypersphere could be equipped with symmetries, of great importance in order to explain central nervous system (CNS) activities, undetectable at the usual 3D cortical level.

Although we live in a 3D world with no immediate perception that 4D space exists at all, the brain hypersphere rotations can be identified through their “cross section” movements on a more accessible 3D surface, as if you recognized some object from its shadow projected on a screen. We may thus evaluate indirect clues of the undetectable fourth dimension, such hypersphere rotations’ hallmarks or signs on a familiar 3D surface. Here we show that, in temporal fMRI series from spontaneous brain activity, MNC discloses the typical patterns of quaternionic rotations and hidden symmetries.

## MATERIALS AND METHODS

### 1 Samples

Spontaneous activity structures of high dimensionality (termed “lag threads”) can be found in the brain, consisting of multiple highly reproducible temporal sequences (Mitra et al., 2015). We retrospectively evaluated video frames showing “lag threads” computed from real BOLD resting state rs-fMRI data in a group of 688 subjects, obtained from the Harvard-MGH Brain Genomics Superstruct Project. We assessed 4 sets of coronal sections (including a total of 54 Images) from the published videos (Threads 1, 2, 3 and 4): http://www.pnas.org/content/suppl/2015/03/24/1503960112.DCSupplemental We favoured studies focused on intrinsic, instead of task-evoked activity, because the former is associated with mental operations that could be attributed to the activity of a torus: “screens” are glued together and the trajectories of thoughts follow the internal surface of a hypersphere. For example, spontaneous brain activity has been associated with day dreaming propensities, construction of coherent mental scenes, autobiographical memories, experiences focused on the future, dreaming state (for a description of the terminology, see (Andrews-Hanna et al., 2014). Each tessellated image leads to the MNC mesh clustering described in the next paragraph.

### 2 Generating Points in Voronoï Tilings of Plane Surfaces

This section introduces nucleus clustering in Voronoï tessellations of plane surfaces (Peters 2016; Edelsbrunner, 2006). A Voronoï tessellation is a tiling of a surface with various shaped convex polygons. Let *E* be a plane surface such as the surface of an fMRI image and let *S* be a set of generating points in *E.* Each such polygon is called a Voronoï region *V*(*s*) of a

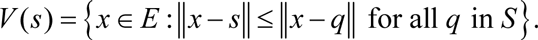

In other words, a Voronoï region *V*(*s*) is the set of all points *x* on the plane surface E that are nearer to the generating point *s* than to any other generating point on the surface (Figures 1A-B). In this investigation of fMRI images, each of the generating points in a particular Voronoï tessellation has a different description. Each description of generating point *s* is defined by the gradient orientation angle of *s*, *i.e.*, the angle of the tangent to the point *s*.

**Figure 1.**
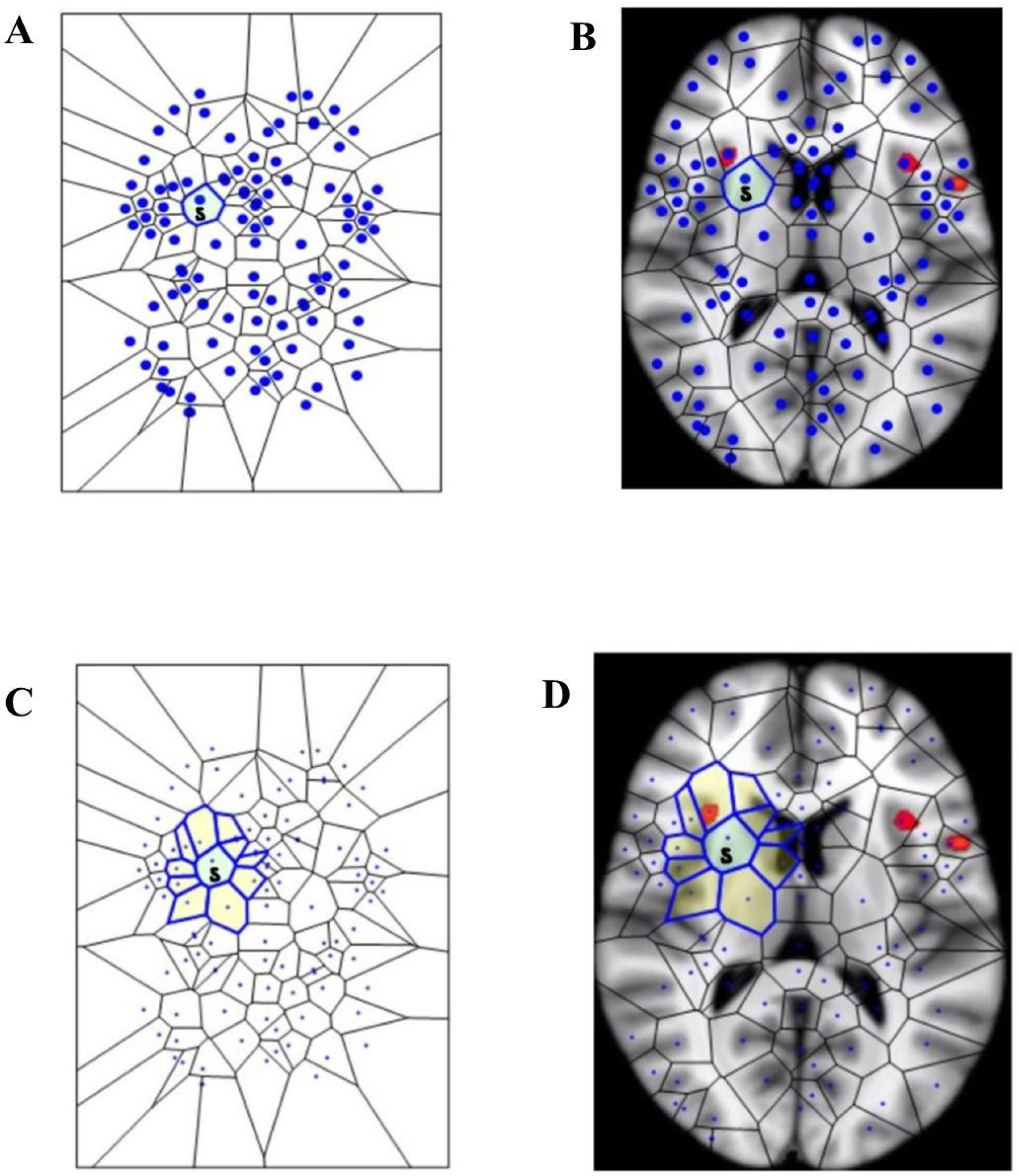
**1A:** surface tiling of a sample Voronoï region 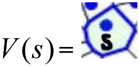. Each •in the tiling represents a generating point with particular features, such gradient orientation and brightness. There are no two • that have the same description. For this reason, every Voronoï region has a slightly different shape. This tiling is derived from the fMRI image in Figure 1B, which displays Voronoï region *V*(*s*) on a fMRI image taken from Mitra (11). Figure 1C displays a sample maximal nucleus cluster *N* for a particular generating point represented by the dot • in N. In this Voronoï tiling, the nucleus N has 10 adjacent (strongly near) polygons. Since N has the highest number of adjacent polygons, it is maximal. This *N* is of particular interest, since the generating point • in N has *a gradient orientation that is different from the gradient orientation of any other generating point in this particular tiling.* In Figure 1D, the maximal nucleus cluster *N* is shown *in situ* in the tiling of an fMRI image.

### 3 Nucleus Clustering in Voronoï Tilings

A *nucleus cluster* in a Voronoï tiling is a collection of polygons that are adjacent to (share an edge with) a central Voronoï region, called the cluster nucleus. In this work, the focus is on maximal nucleus clusters, which highlight singular regions of fMRI images. A pair of Voronoï regions are considered *strongly near*, provided the regions have an edge in common (Peters and Inan, 2016; Peters, 2016). A *maximal nucleus cluster N* contains a nucleus polygon with the highest number of strongly near (adjacent) Voronoï regions (Figures 1CD).

The gradient orientation angle θ of a point (picture element) in an fMRI image is found in the following way. Let *img*(x, y) be a 2D fMRI image. Then

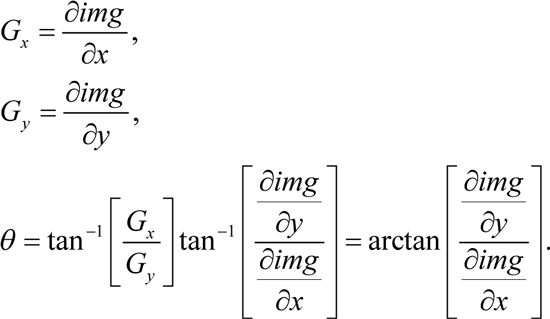

In other words, the angle *θ* of the generating point of mesh nucleus is the arc tangent of the ratio of the partial derivatives of the image function at a particular point (x, y) in an fMRI image.

In sum, for each fMRI temporal frame from Mitra et al. (2015), we produced tessellated images with one or more maximal mesh regions (i.e., a maximal region which contains the maximal number of adjacent regions). Furthermore, we produced tessellated images showing one or more MNC. Each maximal nucleus cluster *N* contains a central Voronoï region - the nucleus - surrounded by adjacent regions, *i.e.*, Voronoï region polygons.

### 4 Steps to Construct a Gradient-Orientation Mesh

Here we give the steps to build Voronoï tiling, so that every generating point has gradient orientation (GO) angle *θ* that is different from the GO angles of each of the other points used in constructing the tiling on an fMRI image (see Figure 2). The focus in this form of Voronoï tiling is on guaranteeing that each nucleus of mesh cluster is derived from a unique generating point. This is accomplished by weeding out all image pixels with non-unique GO angles. The end result is a collection of Voronoï regions that highlight different structures in a tessellated fMRI image. Each Voronoï region *V*(*s*) in a GO mesh is described by feature vector that includes the GO of the generating point *s*.

**Figure 2.**
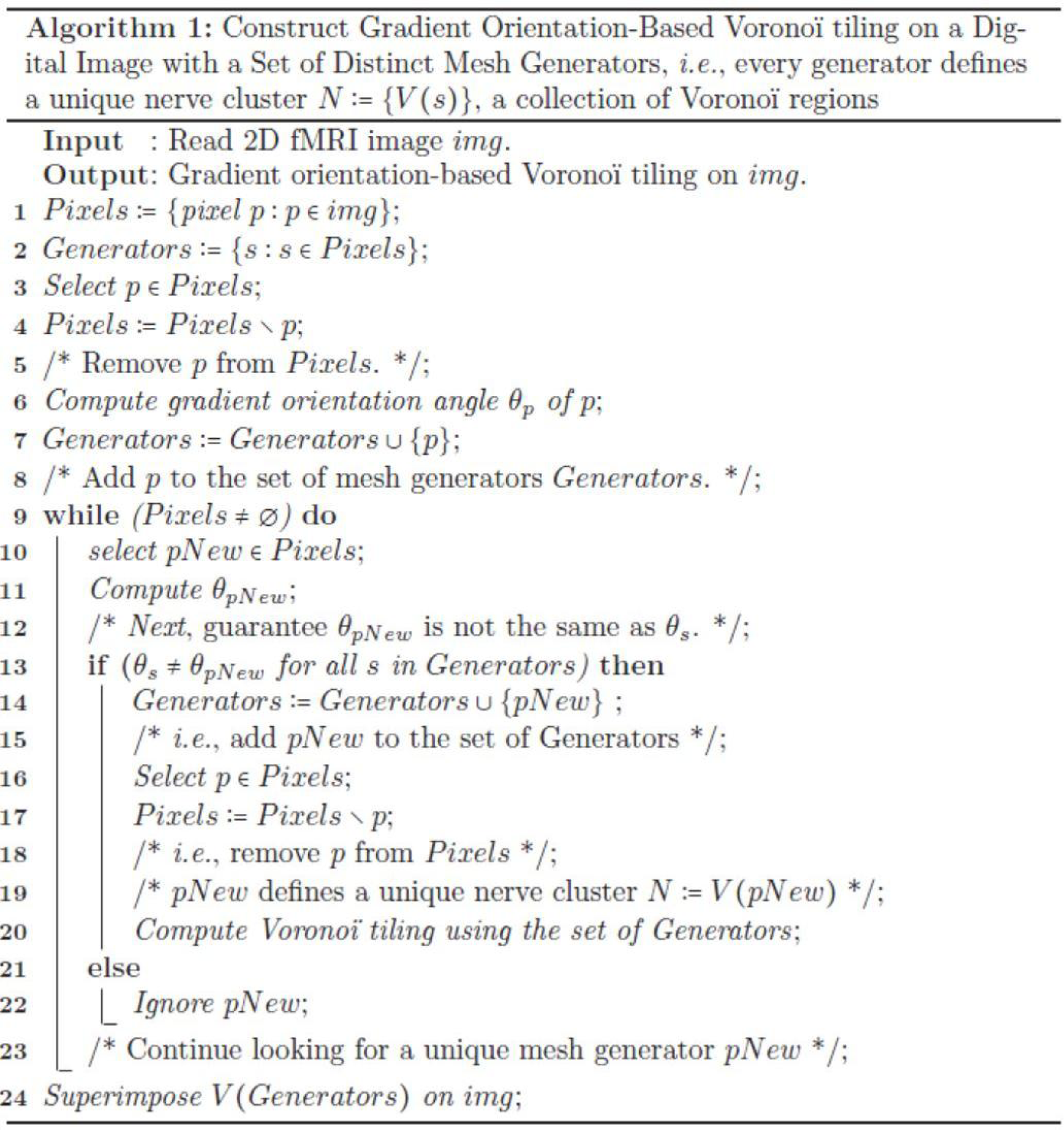
The steps in the method used to construct the mesh on an fMRI image shown in Figure 1D.

Since each s is unique (not repeated in the Generators set in Figure 2), each nucleus mesh cluster *N* has a unique description. Taking this a step further, we identify maximal nucleus clusters on a tessellated fMRI image. In effect, each maximal *N* tells us something different about each region of a tiled fMRI image, since we include, in the description of a maximal nucleus, the number adjacent regions as well as the GO of the nucleus generating point.

### 5 Rényi entropy as a Monotonic Function of Information for fMRI Nucleus Clusters

The major new elements in the evaluation of fMRI images are nucleus clusters, maximal nucleus clusters, strongly near maximal nucleus clusters, convexity structures that occur whenever max nucleus clusters intersect (Peters and Inan, 2016). We showed in the above paragraphs that in a Voronoï tessellation of an fMRI image, of particular interest is the presence of ***maximal nucleus clusters*** (MNC), *i.e.*, clusters with the highest number of adjacent polygons. In this section, we now introduce a measure of the information that MNCs in fMRI images yield. We demonstrate that MNC reveal regions of the brain with higher levels of cortical information in comparison with non-MNC cortical regions, that uniformly yield less information.

In a series of papers, Rényi (Rènyi, 1961; Rènyi, 1966), introduced a measure of information of a set random events. Let *X* be a set random events such as the occurrence of polygonal areas in a Voronoï tessellation and let *β* > 0, *β* ≠ 1, *p*(*x*) the probability of the occurrence of *x* in *X*. Then Rényi entropy *H*_*β*_(*X*) is defined by

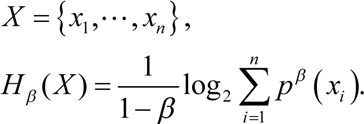

Because of the relationship between Rényi entropy of a set of events and the information represented by events, Rényi entropy and information are interchangeable in practical applications (Rènyi, 1982; Bromiley et al., 2010). In fact, it has been shown that Rényi entropy *H*_*β*_(*X*) is a monotonic function of the information associated with *X.* This means that Rényi entropy can be used as a measure of information for any order *β* > 0 (27).

Let *X_MNC_*, *X_nonMNC_* be sets of MNC polygon areas and non-MNC polygon areas in a random distribution of tessellation polygon areas. Also, let 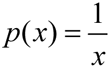, 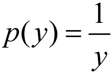 be the probability of occurrence of *x* ∈ *X_MNC_*, *y* ∈ *X_nonMNC_*. Notice that the nuclei in MNCs have the highest concentration of adjacent polygons, compared all non-MNC polygons. Based on measurements of Rényi entropy for MNC vs. non-MNC observations, we have confirmed that Rényi entropy of nucleus polygon clusters is consistently higher than the set of non-MNC polygons (Figures 3 and 4). This finding indicates that MNCs yield higher information than any of the polygon areas outside the MNCs.

**Figure 3.**
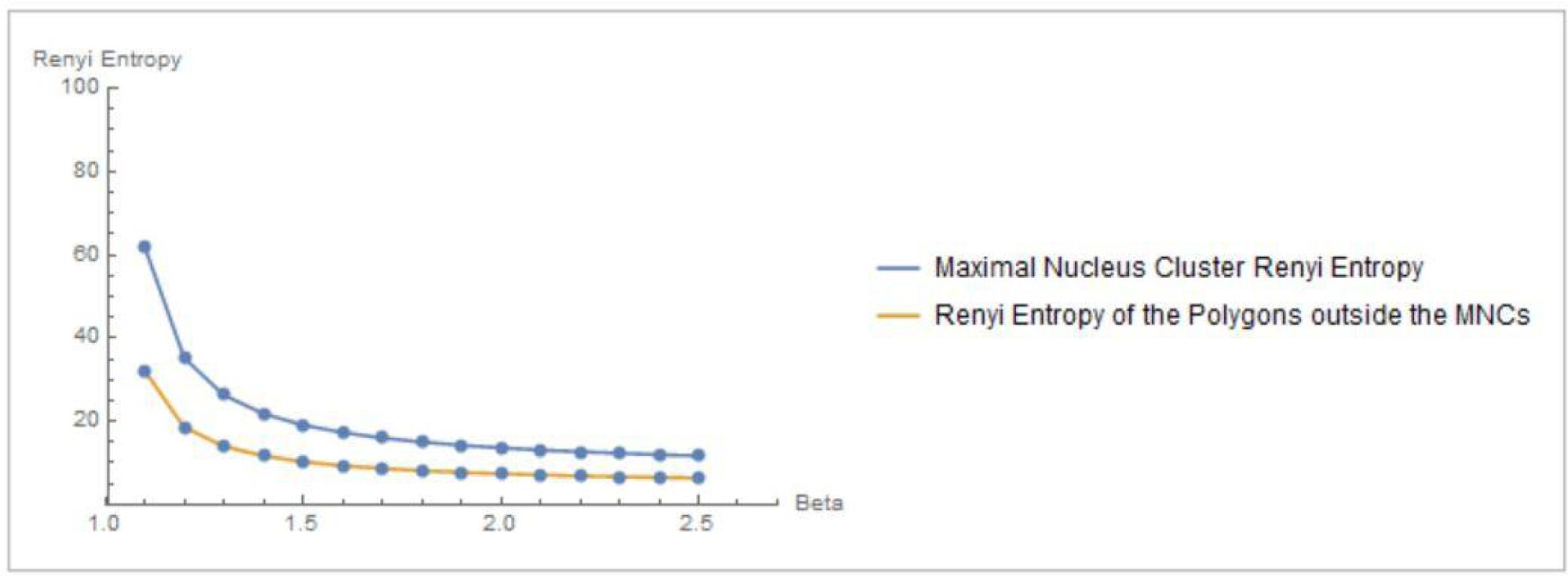
Rényi entropy values of maximal nucleus clusters, compared with the surrounding areas of fMRI images. The *x* axis displays the values of the Beta parameter for 1.1 ≤ *β* ≤ 2.5.

**Figure 4.**
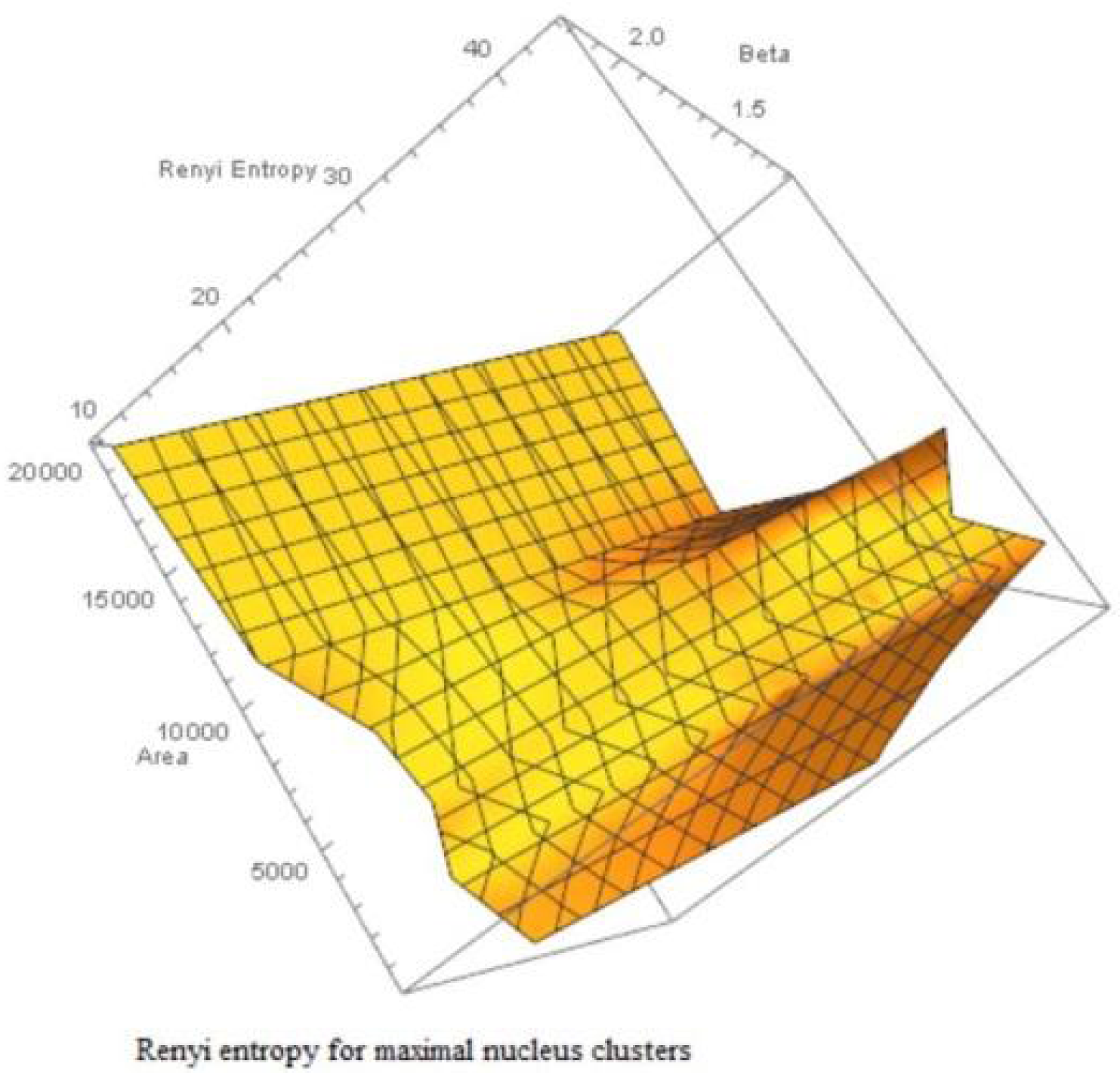
Rényi entropy values vs. number polygon areas vs. 1.1 ≤ *β* ≤ 2.5 of maximal nucleus clusters in fMRI images. MNC Nuclei surround by polygons with smaller areas have higher Rényi entropy, which tells us that smaller MNC areas yield more cortical information than MNCs with larger areas.

In sum, Rényi entropy provides a measure of the information in maximal nucleus clusters and the surrounding zones of fMRI images. This means that the information from areas occupied by MNCs vs. non-MNC areas can be measured and compared. This also means that the maximal nucleus clusters are equipped with higher entropy values (and corresponding higher information), which contrasts with measure of information in the surronding non-MNC zones. Hence, MNCs make it possible to pinpoint the highest source of information in fMRI images.

### 6 Borsuk-Ulam theorem comes into play

The Borsuk-Ulam Theorem (Borsuk, 1933; Dodson and Parker, 1997) states that:

*Every continuous map f: S^n^ → R^n^ must identify a pair of antipodal points (on S^n^)*.

That is, each pair of antipodal points on an *n*-sphere maps to Euclidean space *R*^*n*^ (Beyer and Zardecki, 2004; Matousek, 2003). Points on *S*^*n*^ are *antipodal*, provided they are diametrically opposite (Weisstein, 2015; Marsaglia, 1972). For further details, see Tozzi and Peters (2016a and 2016b). The two antipodal points can be used not only for the description of simple topological points, but also for more complicated structures (Borsuk 1969), such spatial or temporal patterns functions, signals, movements, trajectories and symmetries (Saye and Sethian, 2011; Peters, 2016. If we simply evaluate CNS activity instead of “spatial signals”, BUT leads naturally to the possibility of a region-based, not simply point-based, brain geometry, with many applications (Peters, 2014). We are thus allowed to describe nervous systems functions or shapes as antipodal points on a n-sphere (Figures 3A, 3B). It means that the activities under assessment (in this case, the 4D torus movements) can be found in the feature space derived from the descriptions (feature vectors) in a tessellated fMRI image.

If we map the two antipodal points on a n-1 –sphere, we obtain a single point. The signal shapes’ functions can be compared (Weeks, 2002; Saye and Sethian, 2011): the two antipodal points representing systems features are assessed at one level of observation, while the single point is assessed at a lower level. Although BUT was originally described in terms of a natural number *n* that expresses a structure embedded in a spatial dimension, nevertheless the *n* value can stand for other types of numbers: it can be also cast as an integer, a rational or an irrational number (Tozzi and Peters, 2016b). We might regard functions or shapes as embedded in an *n*-sphere, where *n* stands for a temporal dimension instead of a spatial dimension. This makes it possible to use the *n* parameter as a versatile tool for the description of fMRI brain features (Figure 5C).

**Figure 5.**
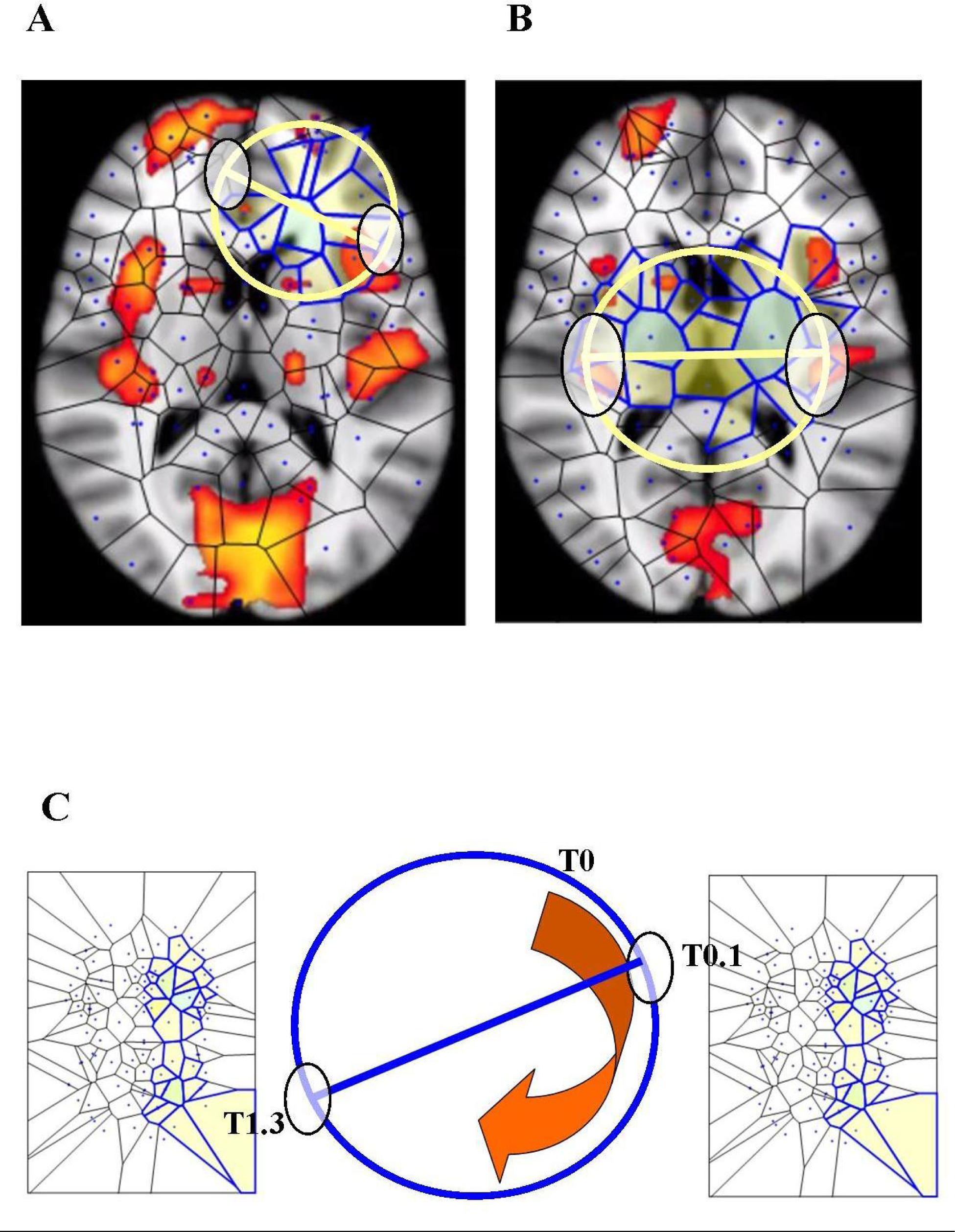
BUT and its variants applied to fMRI neuroimages. **Figures A** and **B** display respectively one and two maximal nuclei clusters ((11), Thread 2). Antipodal points with matching description (on a spatial circumference) can be detected in both the images. Note that the MNC do not necessarily correspond to the “traditional” BOLD activations (shown in red) detectable in fMRI neuroimaging studies. **Figure C** displays a temporal matching description between two maximal nuclei clusters at time T0.1 and T1.3 seconds. Note that, in this Figure, the n-sphere number n refers to the time, and not to a spatial dimension. The curved arrow depicts the time conventionally passing clockwise along the circumference of the n-sphere.

In sum, BUT and its variants say that:

a. There exist regional spatial fMRI patterns (shapes, functions, vectors) equipped with proximities, affine connections, homologies and symmetries.
b. We are allowed to assess the spatial patterns described by the MNC in terms of signals, temporal patterns (in our 4D case, movements and trajectories on the 3D brain), in order to achieve a real-time description of the movements of the hypersphere.

### 7 Quaternionic movements

In a previous study (Tozzia and Peters, 2016a), the presence of a hypersphere was detected invoking BUT: we viewed the antipodal points as brain signals opposite each other on a Clifford torus, *i.e.*, we identified the simultaneous activation of brain antipodal signals as a proof of a perceivable “passing through” of the fourth dimension onto the nervous 3D surface.

Here we evaluated instead, in resting-state fMRI series, a more direct hallmark of the presence of a hypersphere: the trajectory and the temporal evolution of the signals on the 2D brain surface, in order to see whether they match the predicted trajectories of the Clifford torus. To evaluate a hypersphere in terms of a framework for brain activity, we first needed to identify potential brain signal loci where quaternion rotations might take place: we thus embedded the brain in the 3D space of a Clifford torus and looked for its hallmarks or hints (Figure 6).

**Figure 6.**
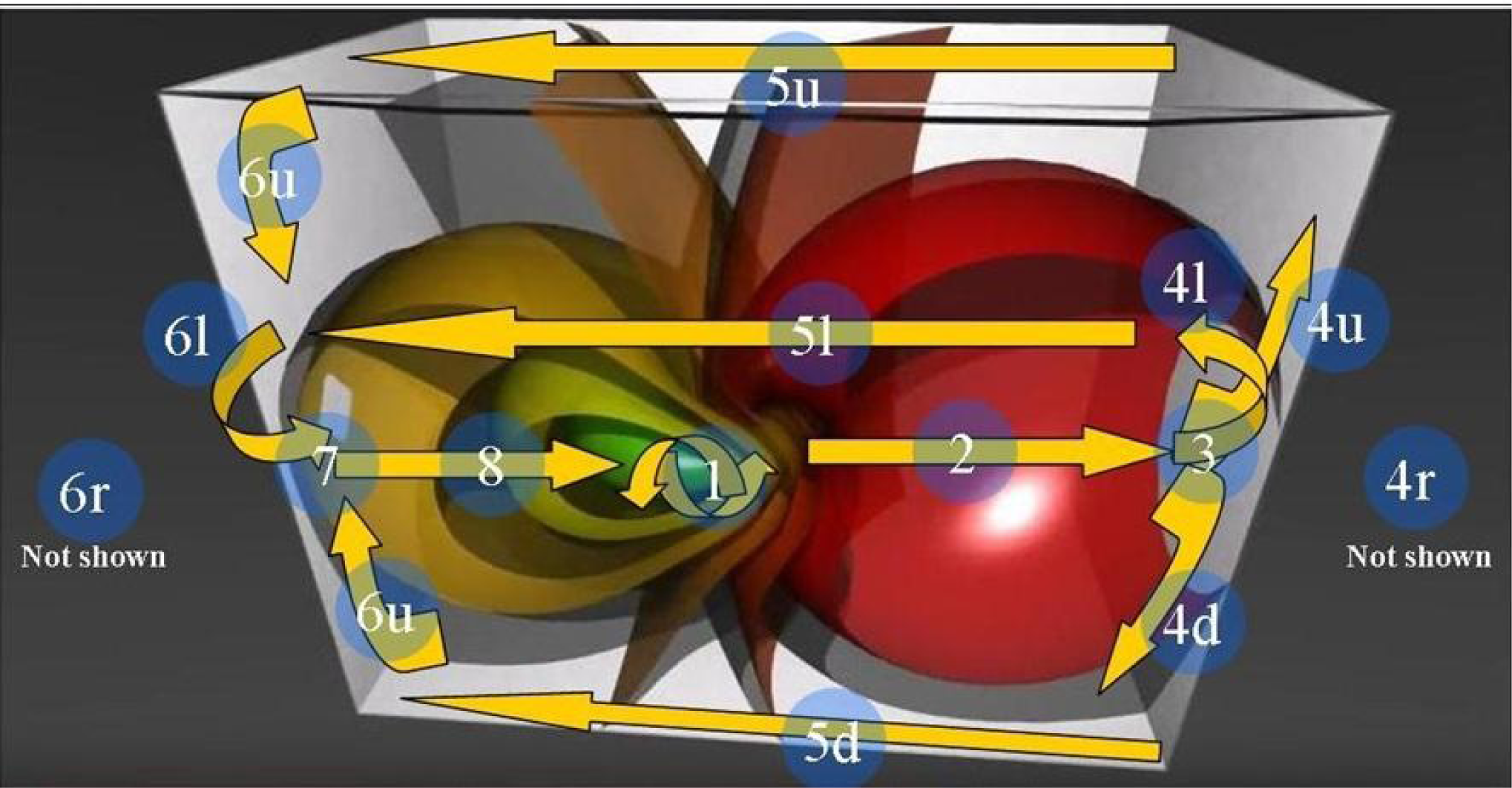
shows the 3D Stereographic projection of the “toroidal parallels” of a hypersphere (from https://www.youtube.com/watch?v=QlcSlTmc0Ts; see also http://www.matematita.it/materiale/index.php?lang=en&p=cat&sc=2,745). The orange arrows illustrate the trajectories followed by the 4D quaternionic movements of a Clifford torus when projected onto the surface of the 3D space in which it is embedded. The circled numbers describe the trajectories, starting from the conventional point 1 (the letters u, d,r, l denote respectively the upper, down, right and left trajectories on the surface of the 3D paprallelepiped). Note that the arrows follow the external and medial surfaces of the 3D space in a way that is predictable. Just one of the possible directions of the quaternion movements is displayed: the flow on a Clifford torus may indeed occur in every plane. In this Figure, the spheres on the right grow in diameter, forming a circle of increasing circumference on the right surface of the 3D space. Conversely, on the opposite left side, the spheres shrink and give rise to a circle of decreasing circumference on the left surface of the 3D space.

## RESULTS

We found that clusters of higher activity, which are equipped with higher Rényi entropy compared with the surrounding zones, are scattered throughout different brain areas. It means that some micro-areas of a specific anatomical zones contain more information than the adjacent ones. In other words, the MNC approach detects zones with higher Rényi entropy, compared with the surrounding ones. In various frames, more than one cluster is detectable. The image data analysis shows also that the MNC activity displays the typical features of the Clifford torus’ movements, supporting the hypothesis that a functional hypersphere occurs during resting state brain activity (Figure 7). The temporal sequence also show the hypersphere moves on the brain, and it moves relatively slowly. At start, the trajectory follows the median sections (see timeframes 0.1-0.3 in Figure 7), then moves towards the posterior part of the brain, where a reflexion of four trajectories along the lateral surfaces occurs (0.4). This pattern closely matches the one predicted by the model illustrating the quaternionic movements on a Clifford torus. The hypersphere does not display a regular or continuous movement, rather it proceeds forward and then backward for a short time (time 0.5 and 0.6). From 0.7, the trajectory follows the patterns predicted by Figure 6.

**Figure 7.**
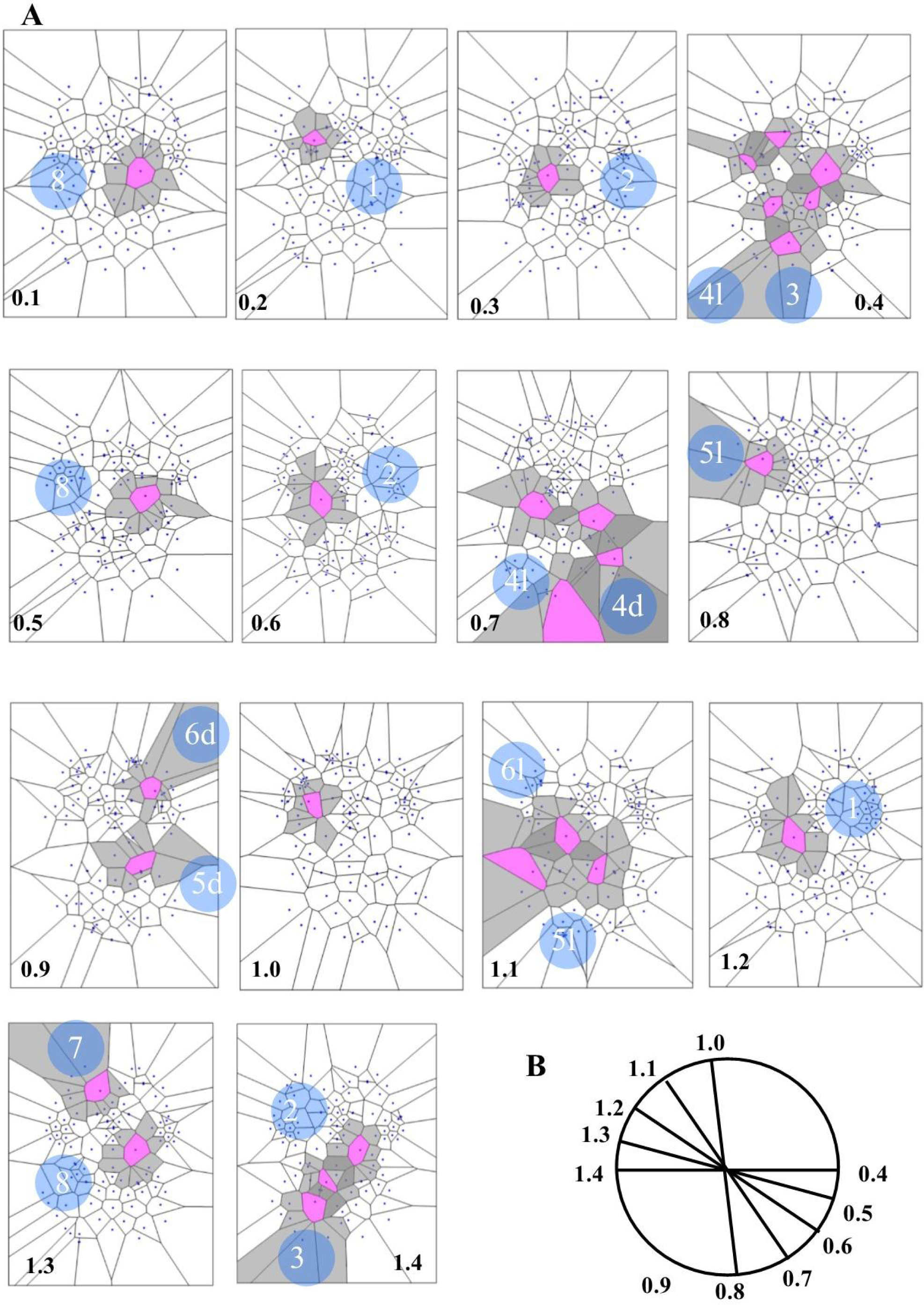
**A** depicts a real pattern of maximal nuclei clusters temporal activation (from Mitra et al., 2015, Thread 4). Note that the typical trajectories of a Clifford torus are clearly displayed (see Figure 6 for comparison). If you look at the parallelepipedal 3D projections of the 4D quaternionic movements (Figure 6), the MNC embedded in the 2D brain surface stand for the 4D movements INTERNAL to the parallelepiped, while the MNC lying outside of the 2D brain surface stand for the 4D movements on the SURFACE of the parallelepiped in Figure 7B. The movements described by maximal nucleus clusters are temporally specular. A matching description among temporal frames occurs, so that, for example, the frame 0.2 displays the same MNC features of 1.3. It means that the hypersphere moves in a stereotyped sequence and according a repetitive temporal sequence, following the trajectories predicted by the quaternionic model.

The MNC activity follows a specular, repetitive temporal pattern of activation (Figures 7, 8, 9). For example, the pictures of the first and the last time have MNC activity with matching description. The MNC activity areas are different from those of BOLD activity (Figure 9). The results can be summarized as follows: the movements of a hypersphere are clearly detectable, and the MNC activity clearly displays antipodal points in a temporal sequence, independent from fMRI activation.

**Figure 8.**
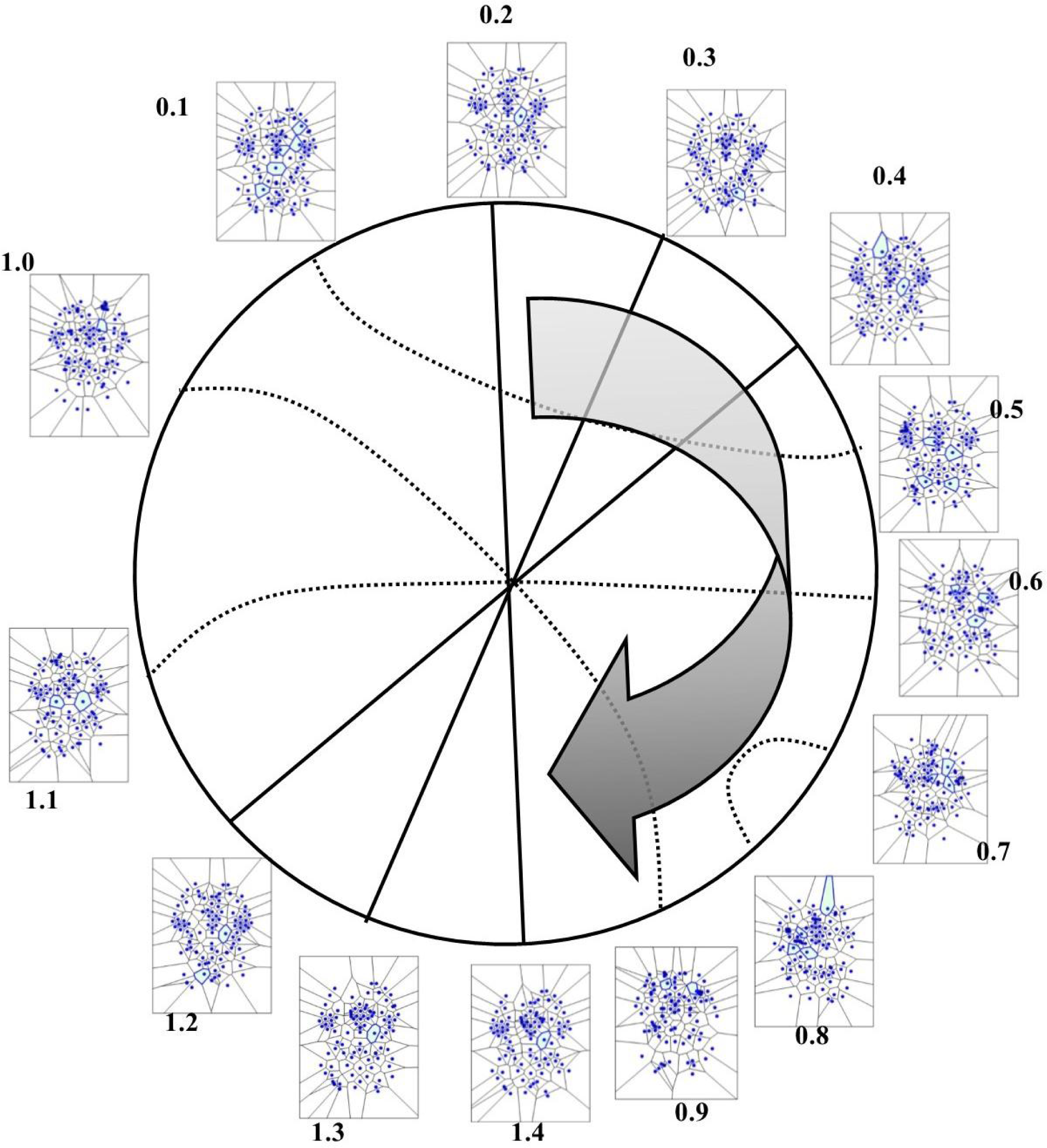
Temporal antipodal points (from Thread 2 frames). The straight lines connecting opposite points on the temporal circumference depict “pure” antipodal points, while the curved lines depict non-antipodal points with matching description.

**Figure 9.**
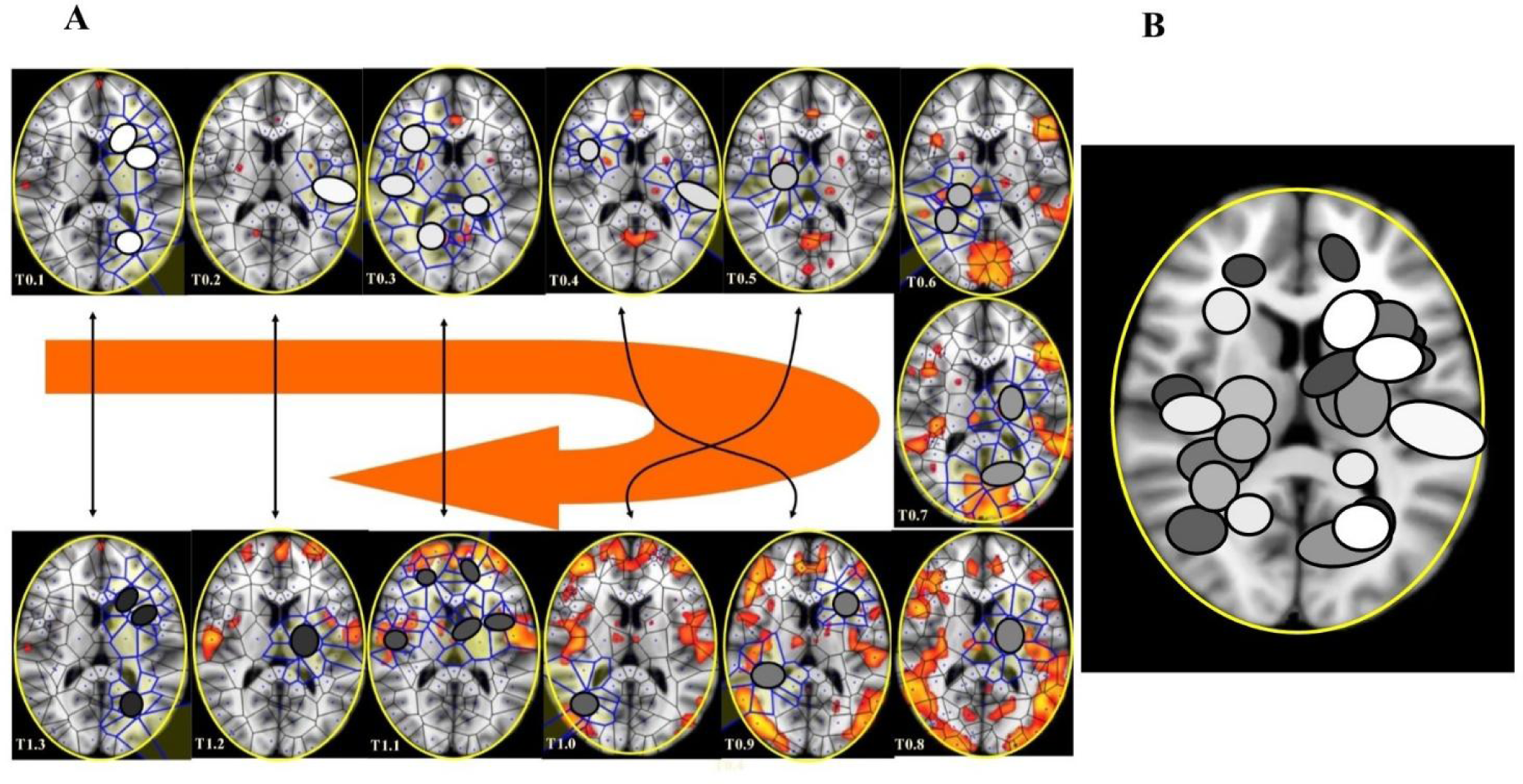
**A.** This figure (from Thread 1 timeframes) illustrates another way to illustrate temporal antipodal points. The temporal sequences are displayed clockwise, from T=0.1 to T=1.3. The Voronoï regions embedded in MNC are depicted as circles. The white circles refer to the presence of mesh clusters at high activity in the initial times, the grey circles in the intermediate times and the darker circles in the later times. Figure 9B. MNC activity on 13 superimposed, consecutive brain images from Thread 1 (from time T0 to T13). Note that the areas of nuclei activation are scarcely superimposed to the “classical” zones of BOLD activation (shown in red).

## CONCLUSIONS

There are two state-of-the-art approaches for understanding the communication among distributed brain systems using fMRI data. The first – dynamic causal modelling - uses models of effective connectivity, while the other - Granger causal modelling - uses models of functional connectivity (Friston, 2009; Friston, 2010). This paper introduces a novel method, the computational proximity, which is different: rather than being correlated with the “classical” BOLD activity, it shows how spatial regions are correlated through their “proximity”. From the experiment done so far, we are beginning to see different forms of brain function represented by the MNC. We showed that computational proximity (i.e., strongly near nucleus mesh clusters) in 2D fMRI images is able to reveal hidden temporal patterns of Rényi entropy, enabling us to detect functional information from morphological data.

Here we have shown that a morphological analysis of simple 2D images taken from fMRI video frame sequences might give insights into the functional structure of neural processing. In a previous study, we evaluated the possible *hints* of a hypersphere on simple fMRI scans during resting state brain activity (Tozzi and Peters, 2016a). We showed how, due to the Borsuk-Ulam theorem (BUT), the fMRI activation of brain antipodal points could be a signature of 4D. The antipodal points predicted by BUT could be evaluated not just on images taken from fMRI studies, but also on datasets from other neurotechniques, such as, for example, EEG. In the current study we used a novel method, in order to confirm the data of the previous work with a completely different and more sophisticated approach. Indeed, looking at the sequences of maximal nucleus clusters and their entropy, we found experimental patterns compatible with the ones predicted by the hypersphere model. We detected on the 3D brain “shadows” of a 4D hypersphere rotating according to quaternion movements: these “hints” make it possible for us the possibility to visualize both the spatial arrangement and the movements of the corresponding Clifford torus. In other words, during spontaneous brain activity, the apparently scattered temporal changes in MNC follow a stereotyped trajectory which can be compared with the 4D movements of a hypersphere. Our study uncovered evidence of hypersphere during spontaneous activity, demonstrating that brain activity lies on a 3-sphere embedded in 4D space. How can be sure that the MNC reveals the presence of a brain hypersphere? Three cues talk in favor of this hypothesis. First, the MNC patterns in resting state fMRI closely resemble the theoretical trajectories predicted by Clifford torus movements. Second, temporal sequences of fMRI images display matching description, in agreement with the BUT dictates. Third, there is a difference in Rényi entropy between MNC and the surrounding zones, these data pointing towards diverse levels of activity. The reproducibility of the hypersphere movements suggests that this organizational feature is essential to normal brain function. Because the Clifford torus incessantly changes its intrinsic structure, due to the different transformations of the quaternionic group, it is reasonable to speculate that each mental state corresponds to a different hypersphere’s topological space. The concept of a hypersphere in the brain is also more noteworthy, if we frame it in the general picture of nervous symmetries (Tozzi and Peters, 2016b). A shift in conceptualizations is evident in a brain theory of broken symmetries based on a hypersphere approach. It might be speculated that symmetries are hidden in a 3D dimension and restored in the higher dimension of the hypersphere, and vice versa. It means that brain functional and anatomical organization may be better assessed if one considers how certain hidden “symmetries”, essential to shaping brain gradients and activity, may only appear under the lens of higher-dimensional neural representations (Tozzi and Peters, 2016b), i.e., the hypersphere. We anticipate our essay to be a starting point for further evaluation, both in physiological and pathological conditions, of a neural fourth spatial dimension where other brain functions, such as perception, memory retrieval, might take place. It is also possible that every brain function displays a peculiar temporal pattern of such a novel entropic “activity”. In sum, we operationalized a novel, fairly inexpensive, image-analysis technique useful in detecting hidden temporal patterns in the brain. We can assess the spatial patterns described by MNC in terms of entropy variations. While, by our “subjective” and “private” viewpoint, we tend to watch an image inferring the semantic parts in order to give it a meaning, MNC allow the detection of the “objective”, entropy, which do not necessarily correspond to the zones of the figure that we repute more significant. This means that MNC provide a basis for quantifying high-yield information areas in fMRI image features that are normally “hidden” from our attention.

Dedicated to the Memory of Som Naimpally

